# Possible Mechanisms Of Action Of Light Inert Gases On Chemiluminescence Arising As A Result Of Lipid Peroxidation

**DOI:** 10.1101/2024.09.12.612290

**Authors:** Iryna Oliynyk

## Abstract

The use of inert gases in biology and medicine and their effect on biological objects both in vitro and in vivo remains an active area of research. It has been established that light noble gases affect antioxidant processes, free radical oxidation, and enhance chemiluminescence, but an explanation of the physical and chemical mechanisms of this effect is still lacking and is key to further theoretical and experimental studies, given the broad prospects for the use of noble gases in medicine. In this article, we present two of the possible mechanisms of light inert gases’ effect on chemiluminescence (CL), a phenomenon that occurs as a result of free radical recombination and chain breakage during lipid peroxidation. Since the effect on oxidation, in turn, precedes the effect on the antioxidant system and the body’s defense mechanisms.

One of the mechanisms of influence is based on the ability of inert gases to dissolve well in lipids and dissolve poorly in water. Their ability to dissolve in lipid bilayers and affect the conformation of lipid complexes can increase the surface area available for oxidation, the surface area that absorbs radiation and reduce the density of the environment, potentially increasing the availability of oxygen for oxidation reactions. This is the so-called spatial mechanism of inert gas influence on oxidation and chemiluminescence.

The second mechanism is based on the influence on the quantum chemical parameters of the reaction medium. The acceleration of VT relaxation processes, the impact on the components of the medium in quenching excited states, and the radiative decay time of the excited state.

## I. INTRODUCTION

Due to their chemical stability and lack of reactivity, noble gases have long been considered biologically neutral. However, modern research demonstrates the significant potential of using noble gases in both medicine and biology. All noble gases have found their application in medicine. However, the most widely used in medicine are the so-called light noble gases: helium, xenon, and argon, which makes them important research objects (see Table 1).

**Table 1.** Disclosure of the mechanisms of inert gas influence on biological systems and explanation of this influence.

| Inert gases | Use in medicine | Research and explanation of physical and biological patterns of action |
| --- | --- | --- |
| He [4] | Radiology[5, 6, 7]<br>Neurology [9, 10, 11]<br>Cardiovascular system [12, 13]<br>Respiratory system [14, 15]<br>Drug delivery [16]<br>Laparoscopy [17, 18]<br>Surgery [19] | Low density makes helium lighter than air, which makes breathing easier [20].<br>Helium quickly removes heat from the body, which can be helpful in treating burns [21].<br>Helium does not dissolve in blood, making it safe for use in respiratory therapy. |
| Xe [22] | Radiology [23]<br>Anaesthesia [24]<br>Neurology [25, 26]<br>Cardiovascular system [27]<br>Respiratory system [28] | Xenon can act as a free radical scavenger, reducing the number of harmful molecules [27]<br>Xenon can regulate the activity of calcium channels in the cell membranes of neurons [29].<br>The chemical shift of Xe changes with time due to the interaction of Xe with liquid molecules. Attractive forces occur between atoms and molecules with induced dipole moments, strong electrostatic attractions occur between hydrogen atoms and electronegative atoms (such as oxygen or nitrogen) [30]. |
| Ar [31] | Neurology [32, 33, 34]<br>Oncology [35]<br>Cardiovascular system [36]<br>Anaesthesia [37]<br>Surgery [38]<br>Ophthalmology [39]<br>Respiratory system [40, 41]<br>Dermatology [42] | Argon inhibits cell proliferation [43]. Argon can limit internal apoptosis [44]. Argon can induce apoptosis and reduce keratinocyte inflammation [45]. Argon can activate antioxidant systems, which can help protect cells from damage caused by free radicals [46]. Changes the expression of cytokines [47] associated with proliferation, apoptosis, inflammation and antioxidant protection. Argon leads to the dissociation of water and the formation of reactive oxygen species [48] |
| Ne | Oncology [49, 50, 51] (used as an ion in radiation therapy) | Heavy inert gases can generate secondary particles like electrons and photons, which can interact with tissues. Similarly, heavy inert gases can generate delta rays, densely ionizing tracks of secondary electrons that cause severe DNA damage [52]. |
| Kr | Anesthetic effects [53] | Inert gases (Xe, Kr) can deactivate proteins by sticking their molecules together. This reversible process causes proteins to lose their shape and functionality. The study showed that Xe and Kr deactivate pepsin (a digestive enzyme) depending on gas concentration and temperature. Possible mechanism: inert gases accumulate near the active areas of the protein, disrupting its structure and binding to the substrate [54]. It is important to note that the effects of inert geysers depend on a number of factors, including concentration, exposure time, and type of mixture. Some studies have shown that inert gases can have a toxic effect on cells, or have no effect at all [55]. |

Noble or inert gases share common characteristics, including: chemical inertness, low solubility in water, and the absence of an electrical charge 1. This is due to the fullness of the chemical shells. This implies safety for use. At the same time, scientific research is constantly providing information on the biological activity of noble gases and their interaction with biological systems. Modern research shows that noble gases can exhibit anti-inflammatory, anesthetic, analgesic, and neuroprotective properties. These properties make them promising candidates for use in medicine, in particular for the treatment of neurodegenerative diseases, coronary heart disease and other pathological condition.

In addition, without directly participating in the oxidation reaction, the authors established the influence of inert gases on the antioxidant system 2. It is not correct to call inert gases pro- or antioxidants, since antioxidants inhibit oxidation, and inert gases can most likely be called synergistic products, and thus determine their ability to influence the processes of redox reactions. Thus, inert gases, by affecting oxidation processes, have the ability to protect cells and tissues from damage caused by reactive oxygen species (ROS).

In this case, inert gases correct redox processes and, accordingly, reduce the negative effects of ROS, enhancing antioxidant protection. But again, we remind you that argon increases the intensity of chemiluminescence, which is a consequence of radical recombination. So how do inert gases, by increasing radical recombination, help prevent oxidative stress, which is one of the key causes of cellular damage?

Since inert gases are the cause of antioxidant protection without stimulating free-radical oxidation, they can be safely used in the therapy of diseases where oxidative stress plays a central role, such as in neurodegenerative conditions or ischemic heart muscle damage [3].

Tab. 1. demonstrates not only the variety of inert gases used in medicine, but also contains possible explanations for these effects. The most widely used inert gases in medicine are helium and argon. This is primarily due not only to the medical and biological effects, but also to the cost of these noble gases.

The prevalence of argon in nature and its ease of retention not only makes it used in a wide range of medical applications from surgical procedures to diagnostics, but it is also included in the mixture of scuba gear.

Although more expensive than argon, helium is an important component in medical technology, in particular in the cooling systems of magnetic resonance imaging (MRI) scanners, and is automatically used for burns because of its high thermal conductivity, and is also included in breathing mixtures for patients with respiratory disorders. The complexity of its extraction and low abundance in the atmosphere makes helium much more expensive than argon.

Neon, Krypton, and Xenon are used less frequently not because of their medical and biological effects, but because of their high cost, which is a direct consequence of the difficulty of extracting these gases from the air and their low concentrations in the atmosphere. Therefore, despite the fact that Xenon has good anesthetic properties, the cost limits its widespread use, making it available only to specialized medical centers and research programs.

The table shows that, despite the fact that noble gases demonstrate a large number of biomedical effects, physicochemical explanations for these effects are insufficient. Sometimes, the assumptions for explaining the effects of noble gases are not correct due to the physicochemical characteristics of noble gases.

For example, the fact that argon can accelerate the oxidation of chloride by hydrogen peroxide in aqueous solution, which is manifested in the enhancement of luminol-dependent CL [48], is explained by the fact that argon can affect the water base of biological systems, increasing water dissociation and the formation of reactive oxygen species [48]. However, these assumptions are incorrect, because argon, like other noble gases, is characterized by extremely low (actually absent at physiological temperatures) chemical activity and very low solubility in water and blood plasma. The filled outer electron shell makes it practically incapable of forming chemical bonds or participating in chemical reactions.

Since the destruction of covalent bonds in a water molecule requires a significant amount of energy, light inert gases and argon in particular do not have enough energy to initiate this process. In addition, there are no reliable experimental data confirming that the dissolution of argon in water leads to a significant increase in water dissociation and the formation of reactive oxygen species. But the fact that argon enhances the intensity of chemiluminescence remains. If we add the effects of chemiluminescence enhancement to the effects on the antioxidant and free radical system, there is currently no explanation for these effects in the literature. The present study aims to demonstrate two possible effects of light inert gases such as helium, xenon and argon on the chemiluminescence of aqueous solutions and on the antioxidant system, and to explain possible neuroprotective and cardioprotective effects, which are often accompanied by rather controversial results.

## II. RESULTS

Chemiluminescence (CL) is a superheat radiation. In biofluids, it is a phenomenon of lipid peroxidation reactions that occur as a result of chemical reactions with oxygen. And as a result of the recombination of peroxide radicals, excess energy is released in the form of light quanta in the visible and near-infrared regions of the spectrum. [56, 57].

According to modern concepts, many vital metabolic and physiological processes occurring in the body are closely related to lipid peroxidation (LPO). LPO takes place in the following stages: initiation of the radical chain, continuation of the radical chain, and branching of the radical chain [58]. A chain break leads to a loss of an unpaired electron.

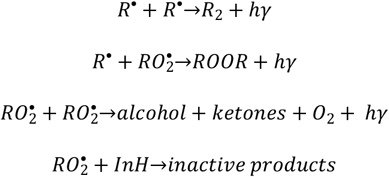

As you can see, at the last stage, the radical breaks with the emission of light quanta. This break with light emission is defined as biochemiluminescence. Initiators of oxidation include active forms of oxygen, metals of variable valence, irradiation, and oxidants. Antioxidants, as well as lack and excess of oxygen gas molecules, act as regulators. The amount of light produced by CL is proportional to the amount of free radicals present. Therefore, CL is often used as a quantitative indicator of free radical oxidation.

In medical diagnostics, CL is used to detect the presence of free radicals in biological samples. This can be used to diagnose diseases such as cancer and heart disease [59], and will also allow us to understand the possible influence of light inert gases on oxidation processes.

Because of CL carries information about the intensity and mechanism of free-radical, non-enzymatic lipid oxidation, let’s explore the mechanism of CL in the body. In contrast to oxidative reactions that occur in vitro, in vivo reactions occur in the environment inside the body. This type of environment represents a dispersed system, in which reactive oxygen species (ROS), denoted by *O*^•^, readily react with lipids (L), which are already present in complexes before the interaction (Fig 1). These lipid complexes (*LCs*) are generally inert and consist of a large number of tightly packed, stable molecules. Interestingly, despite their inert nature, *LCs* can still participate in many oxidation processes.

**Fig. 1.**
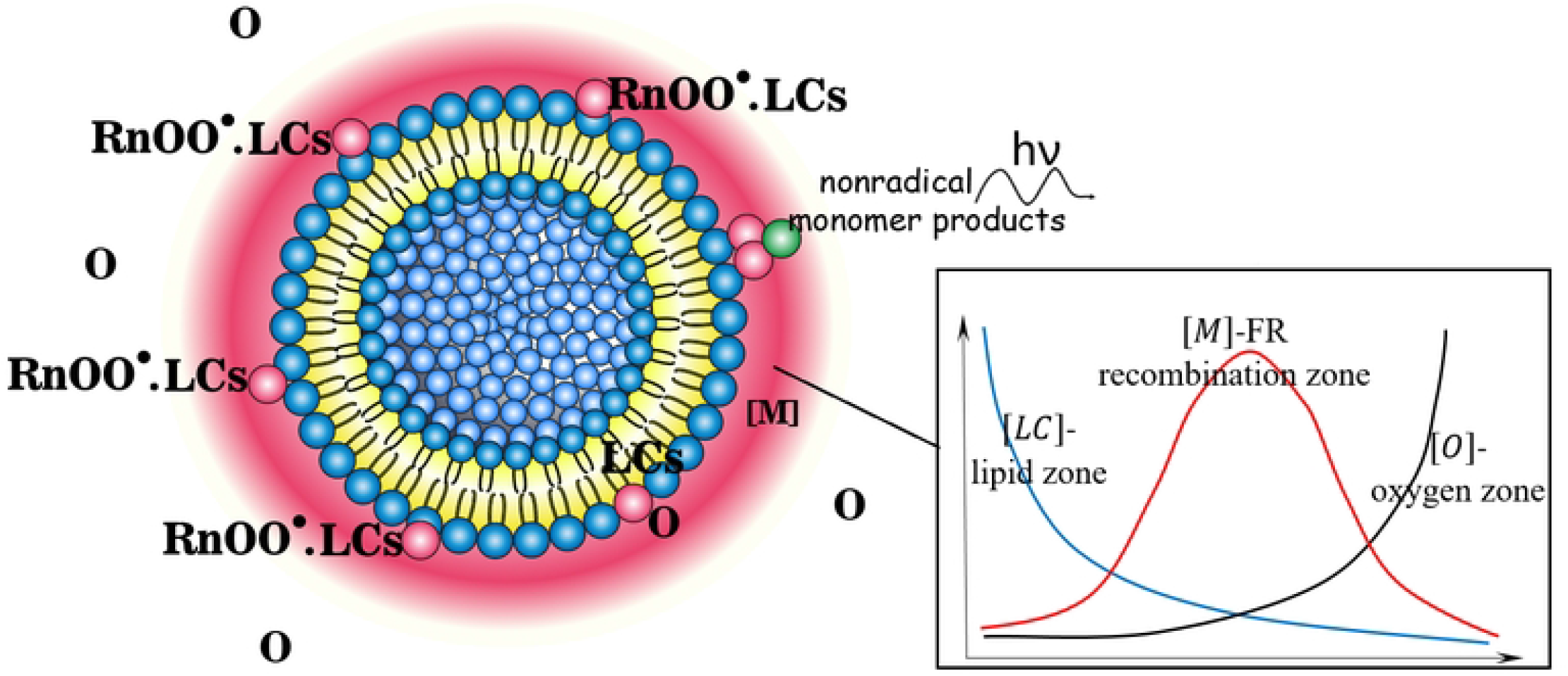
The oxidation reaction, occurring under steady-state conditions with a random lipid from the cluster and leads to chemiluminescence

The figure schematically demonstrates the mechanism of oxidation of lipid complexes. Note that lipid complexes can consist not only of a double layer of lipids, but also of a monolayer of lipids. Oxygen passes through the medium [*M*] and only after reaching the lipids can oxidation occur, the denser the layer of the medium, the more difficult it is for oxygen to penetrate the lipids. Medium [*M*] consists of free radical oxidation products and oxygen, and it is in this layer that free radicals recombine. And it is this layer that is responsible for the glow. In addition, the graph tab schematically shows the zoning of element concentrations depending on the distance to the membrane surface. Three zones are schematically distinguished: the lipid zone (blue line), the recombination zone (red line), and the oxygen zone (black line). This comprehensive schematic illustration demonstrates how radicals, light, and oxygen can interact with these lipid components to affect the structure or stability of membranes.

The oxidation reaction, occurring under steady-state conditions with a random lipid from the cluster, does not affect the overall properties of the system. This allows the lipid complex to participate in multiple processes. Oxidation processes in such a system are likely to proceed with a higher probability due to the increased collision duration. LPO and radical recombination, in turn, lead to CL. Let’s schematically depict the process taking place:

1. Initiation (formation of ab initio lipid free radical)
2. Propagation

Free radical chain reaction established

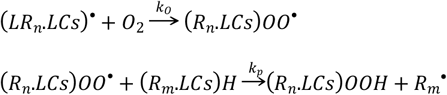

Free radical chain branching (initiation of new chains)

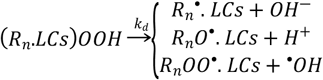

III.Termination (formation of non-radical products)

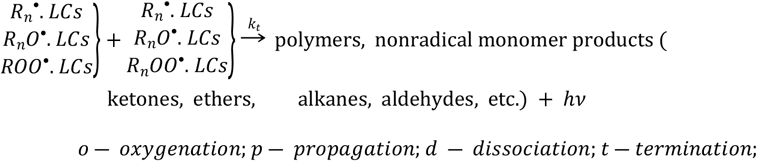

in accordance with the above *LC* = ∑_*j*_ *L*_*j*_, *j* ≫ 1

The final phase of lipid peroxidation (LP) generates highly energetic (transient) entities capable of either returning to a stable state or releasing excess energy in the form of emitted light, which consequently constitutes a byproduct of LP. For example singlet oxygen, ^1^*O*_2_, which can transition back to a stable triplet state, emitting faint phosphorescence in the near infrared spectrum around 1270 nm. This emission (CL), has been extensively utilized to monitor the production of ^1^*O*_2_ during LP and various methods have been devised to amplify CL in biological systems for heightened detection sensitivity [60].

In addition, chemiluminescence emission in the visible range is also observed during LP, which is primarily associated with the recombination of excited carbonyl compounds in the triplet state. These include those formed in the Russell termination mechanism.

The intensity of CL is expressed according to [61]:

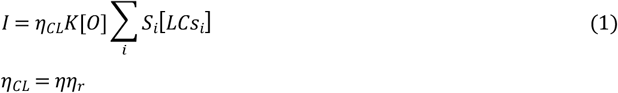

where:

*η*_*CL*_–quantum yield of CL,

*η* –the probability of the formation of an excited molecule upon collision with a lipid complex, or the probability of excitation of an emitting level (*η* should not depend on the size of the clusters), *η*_*r*_– the probability of light emission by an excited molecule.

Additionally that excited states decay over time *τ*_*r*_ with the emission of light and are extinguished, with the constant of the extinguishing rate *k*_*q,i*_ on the components of the solvent medium *M*_*i*_, meaning:

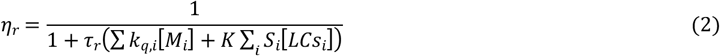

where *S*_*i*_– cross section of fragments of the same size *i*, [*LCs*_*i*_]– concentration of fragments of the same size *i*.

In accordance with the above, the intensity of CL will be written in the form:

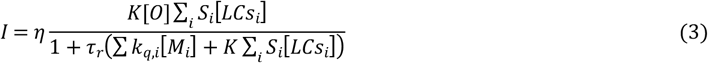

where *K* – the average rate of oxidation reactions.

In order to more thoroughly explain the influence of inert gases on oxidative processes, it is necessary to determine the conditions for the generation of glow by excited molecules.

In CL reactions, a group of electronic states of the molecule, including the ground state, are usually populated.

A characteristic feature of LPO is the population of the excited and ground states of emitter molecules (EM) during the termination (III) reaction process. These excited molecules can then emit light. EM, also participates in the conversion of vibrational energy into translational energy (*VT* - relaxation). In the LPO process, environment molecules can promote *VT* -relaxation of excited EM.

The deactivation of the excited state of a molecule can occur in a variety of ways, including light emission, chemical reaction, and *VT* -relaxation. In the case of EM, which are chemically unstable molecules, deactivation usually occurs by chemical reaction or *VT* -relaxation.

During the deactivation processes of the excited state *g*′, a group of vibrational levels 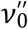 is populated, which is located above the levels on which phototransitions end *g*′*v*′→ *g*′′*v*′′.

EM are the final product of the LPO process. However, there are significantly more EM in the ground state than in the excited state. Therefore, for a time that is longer than the decay time of the excited state of EM, the amount of product that actually causes luminescence in the ground state is greater than in the excited state. This can be demonstrated in particular by the typical curves of initiated CL, which reaches a plateau of stationary luminescence.

Consider schematically the transformation of the luminescent product, during excitation and depletion of levels, see Fig. 2.

**Fig. 2.**
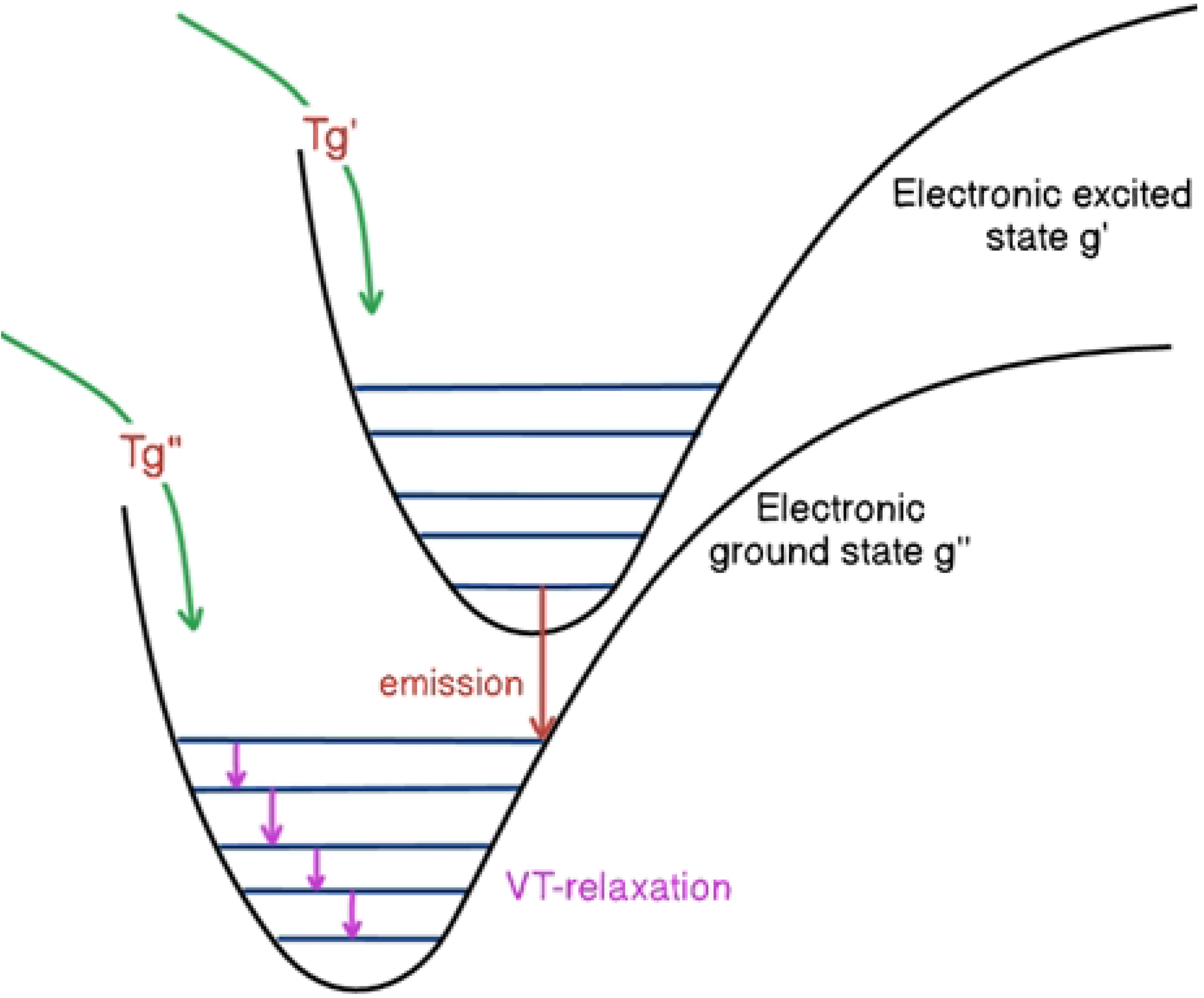
The scheme of a chemical reaction between two electronic states of molecules with product accumulation: curved arrows indicate the directions of chemical transformations, straight arrows - emissive and non-emissive transitions, when the molecule dissipates energy through interaction with other molecules.

Let the consequence of the chemical reaction be the settlement of the basic *g*′′ and excited *g*′ electronic states. The condition of population of the main state is (full electronic inversion)

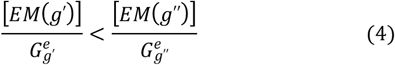

where [*EM*(*g*′)] and [*EM*(*g*′′)] EM (product CL) in the excited and ground states, respectively, and 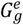-is the multiplicity of degeneracy of the corresponding electronic state. Note that unlike fluorophores, CL products can participate in light emission only once. Further, only after chemical transformations, they can again turn into excited molecules capable of recombination by light radiation.

The concentration at an arbitrary vibrational level (*v*) is determined (using a quasi-stationary distribution over vibrational levels): [62]:

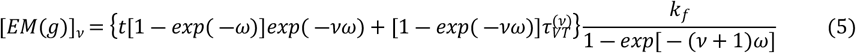

Where *k*_*f*_-is the rate constant of the reaction that leads to the formation of EM, *t* is the reaction time, *ω* ≡ *hω*_*ev*_ *kT, T*–is the temperature of the environment, and 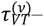 is the vibrational relaxation time of *g, v* levels.

Using the geometric progression formula 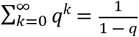 we get

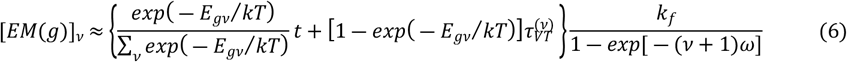

where *E*_*gv*_ is the state energy determined by the set of quantum numbers.

The last multiplier represents the number of chemical transformation (termination) events per unit volume per unit time that lead to the formation of EM in both the ground and excited states, denoted as 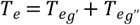.

Considering that for time intervals 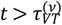, the majority of molecules reside in lower energy levels, we can express 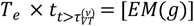.

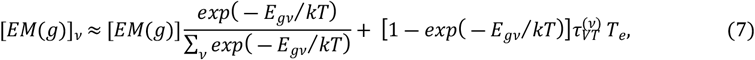

For the excited state *g*′, the *VT* -relaxation time is very small compared to the *VT* -relaxation time of the state *g*′′. Therefore:

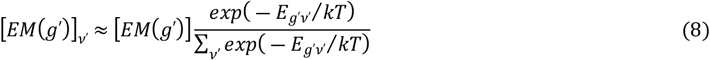

Considering that the excited state *g*′ decays radiatively with a time *τ*_*r*_ and is quenched with a constant quenching rate *k*_*q,i*_ on the components of our medium *M*_*i*_, the concentration of molecules in the state *g*′ is given by:

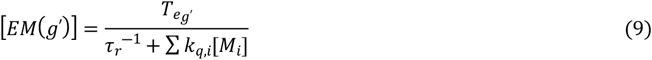

Therefore, equation (8) can be rewritten as:

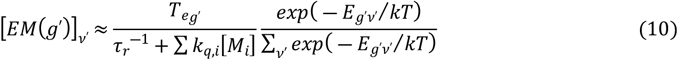

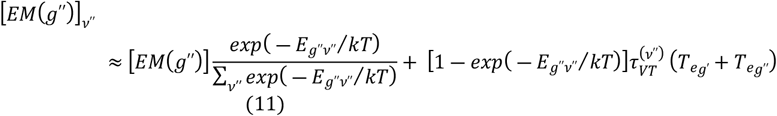

For radiation to exist, a critical condition known as the inversion population criterion must be met. This criterion, specifically for lines with the strongest intensity in the vibrational band *g*′*v*′→ *g*′′*v*′′, is expressed as:

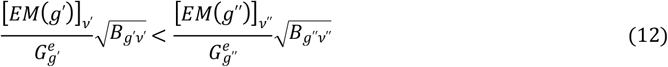

where:

[*EM*(*g*′)]_*v*_′: Population density of molecules in the excited state *g*′ at vibrational level *v*′

[*EM*(*g*′′)]_*v*_′′: Population density of molecules in the lower state *g*′′ at vibrational level *v*′′

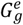: Electronic degeneracy of state ***g***

*B*_*gv*_: Rotational constant for state *g* and level *v*

This inequality implies that the population density in the excited state must be greater than that in the lower state for radiation to occur.

Substituting Population Densities: by substituting the expressions for [*EM*(*g*′)]_*v*_′ and [*EM*(*g*′′)]_*v*_′′ based on the rate equations, we obtain:

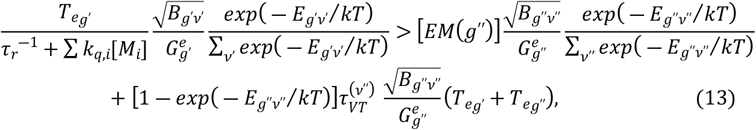

Let’s consider cases of fulfillment of this inequality.

To simplify, let the last term of the right-hand side be the main contributor to the inequality.

Let’s enter the probability of excitation of the level from which radiation occurs:

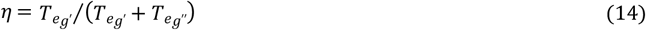

Next, we introduce the generalized rate constants:

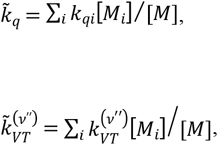

and given that 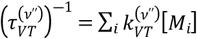, and 1 ― *exp*(― *E*_*g*′′*v*′′_ /*kT*) ≈ 1 (the energy of the state is much greater than the thermal energy of the movement of liquid molecules), we obtain after transform the inequality:

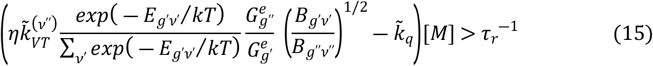

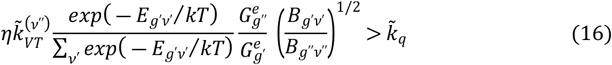

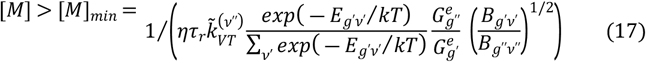

Suppose for working levels *v*′′, the following condition holds:

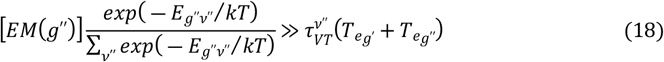

In other words, if the first term dominates, then we obtain another necessary inequality:

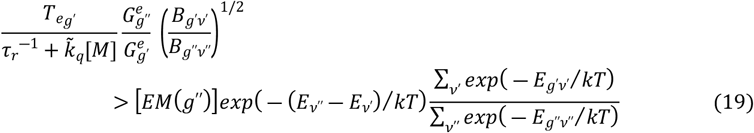

## III. DISCUSSION

Thus, we have come to two mechanisms of inert gases’ influence on lipid peroxidation. The first mechanism is based on the ability of inert gases to penetrate lipid complexes, increasing the absorption of radiation on the molecules of the complexes, and the second mechanism is responsible for the quantum-chemical conditions of radiation existence. Let’s analyze the results in more detail and see how they can be consistent with the experimental data. Note that only the first mechanism can directly affect oxidation.

EM are end products of lipid peroxidation. EM are excited molecules that decay easily. They decay by radiation and are the final products of chemical reactions. High numbers of excited molecules can be harmful to the body because they can damage cells. Excited molecules possess more energy than those in their ground state. This excess energy can be transferred to other molecules, causing damage [63].

For example, EMs can transfer their energy to DNA molecules [64], which can lead to mutations. They can also transfer their energy to protein molecules, which can lead to their degradation and protein transformations [60].

Grounding excited EMs is beneficial because it reduces the risk of them damaging cells. When excited EM emits light, it loses its energy. This makes it less likely that energy will be transferred to another molecule and free radical reactions will continue to branch.

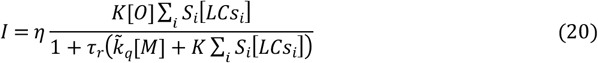

It can be seen from (20) that the ratio of concentrations of reagents in the medium strongly affects the intensity of the glow. The environment itself also contributes to the process. The stronger the diluted solution 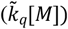, the less quenching on the molecules of the medium, and accordingly, the increase in the level of CL should be observed we will observe. The intensity increases with increasing oxidant concentration [*O*], as well as the total cross-section of lipid complexes.

That is, CL is not only a fact of LPO, but also a natural protective phenomenon in dyslipidemias. The effect on CL reflects the effect on one of the links of metabolism. As a result of a sharp increase in the concentration of the oxidant or substrate in the reacting volume, the intensity of CL should increase (20). Severe injuries, which dramatically change the reaction environment in a certain volume and area of damage, can significantly affect the intensity of CL. In particular, leading to sharp growth ∑_*i*_ *S*_*i*_[*LC*_*i*_].

From the above (20), the effect of inert gases can be on increasing the size of lipid complexes that occur after dyslipidemia. This is due to the inertness of atoms of inert gases, their poor solubility in water, and good solubility in lipids. Inert gases integrating into lipid complexes do not actually increase the numerator, since they do not increase the number of lipids capable of oxidation, but they do affect the term in the denominator, ∑_*i*_ *S*_*i*_[*LC*_*i*_], which cannot be arbitrarily large, since it determines the inverse path length of atoms with respect to capture on fragments:

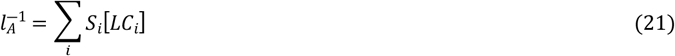

It has already been emphasized that inert gases lead to a dilution of the reaction zone [*M*], the density of which will decrease. This, in turn, will lead to a decrease in the denominator of the term responsible for quenching the excited state on the molecules of the medium 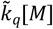. This, in turn, will also lead to an increase in the intensity of CL because a compensatory mechanism will occur - an increase ∑_*i*_ *S*_*i*_[*LC*_*i*_] and a decrease in 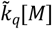. And depending on the initial area of the lipid complexes and the integration of inert gases into them, we can have different effects.

As a result of prolonged exposure to the inert gas, equilibrium will eventually be established.

The first mechanism is directly related to the second mechanism. Since the existence of luminescence requires the recombination of free radicals, which leads to the existence of products in an excited state. In this case, noble gases have an impact on quantum chemical processes. In order for the glow to exist in the working zone, it is necessary also a must to fulfill the inequalities (15-19). It is these ratios that determine the heterogeneity of the results of experiments and the influence of inert gases in various diseases, and help to understand why the influence of inert gases is sometimes even contradictory to the scientific literature, since the obtained ratios impose conditions on the medium, kinetic parameters of chemical reactions, and on the molecules of the CL emitters themselves. Because the formation rate of EM in both ground and excited states is solely determined by their concentration, these inequalities effectively translate to requirements for the concentrations of reactants, products, and the overall mixture composition that influence the quenching rate.

The first thing to notice in these ratios is the temperature. Lowering the temperature improves the performance of the condition (19), which actually proves why in a significantly affected organism with high sudden hyperlipidemias, lowering the temperature has a significant therapeutic effect. But a significant decrease in temperature can violate other strict inequalities.

Because 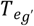-this is the rate of appearance of EM in the excited state, it clearly depends on the concentrations of reagents, in particular, oxygen entering the reaction zone and the cross-sectional area of lipid complexes, as well as the rate of quenching of excited states that participate in oxidation (19). Vibrational relaxation is not clearly included either in equation (3) or in the final inequality (19), but inequalities (15, 16) impose a condition on the rate of *VT*-relaxation. Equation (16) reveals a distinct dependency between the rates of vibrational relaxation (*VT*-relaxation) and electronic quenching of the excited EM within the medium’s components. Crucially, *VT*-relaxation must be considerably faster than quenching (transition to the ground state). Accordingly, with hyperlipidemia, which occurs with a large number of pathologies, excited products of radical recombination, such as nonradical monomer products (ketones, ethers, alkanes, aldehydes, etc.) quickly transition to the ground state, and do not lose energy due to quenching on the components environment, the presence of a component that weakly quenches the emitter molecule, but effectively participates in relaxation processes, is necessary in the reaction zone. These components are the light inert gases *He, Ar*. Since they are poorly soluble molecules in water, they can easily penetrate into the reaction zone and serve as a solvent that does not take part in chemical reactions, and they also take an active part in *VT* - processes [65]. At the same time, the active zone of the reaction cannot be diluted too much, this follows from inequality (17). In particular, inequality (17) indicates the limitation of the lower limit of the concentration of deactivator molecules (substrate molecules). The minimum concentration of molecules and subsequent substrate dilution is influenced by the speed of *VT*-relaxation. A faster deactivation of lower energy levels (*g*′′ *v*′′) is necessary compared to the rate of radiative decay from the state *g*′.

This means that at low concentrations of the oxidizing substrate (weak lesions), inhalation of inert gases may not have ЯВнОГО therapeutic effect. Additionally, it indicates the limitation of the therapeutic duration of inhalation in such cases. Prolonged use of light inert gases will increase the rate of *VT*-relaxation but decrease the substrate density in the reaction zone, leading to a decrease in *τ*_*r*_ and necessitating an increase in the medium density. Therefore, inert gases can have a rapid therapeutic effect only in cases of sufficiently severe diffuse or focal lesions characterized by significant hyperlipidemia and increased average lipid density in the lesion. In lesions without hyperlipidemia, the duration of inert gas action must be shortened to prevent excessive medium dilution. Under appropriate conditions, inert gases efficiently utilize the energy of free radical oxidation. Since inert gases do not participate in chemical transformations, their impact on the oxidation processes of the rest of the body, not meeting the outlined conditions, will be insignificant. The significant variation in the therapeutic effects of light inert gases in various articles is primarily due to the oversight of hyperlipidemia in the affected area. Some authors highlight a substantial impact and effectiveness from their use, while others report no effect. This discrepancy is influenced by secondary oxidation processes, emitter deactivation on medium molecules, extensive medium dilution, and the duration of exposure to light inert gases.

## VI. CONCLUSIONS

Thus, despite their inertness, lack of charge and low atomic mass, light inert gases are able to influence both free radical oxidation and enhance chemiluminescence, and, accordingly, to influence oxidative stress, which occurs as a result of the accumulation of free radicals in cells and leads to harmful effects on cell membranes, proteins and DNA. Inert gases do not affect the oxidation mechanisms at the chemical level, but affect the spatial and temporal distribution of products involved in free radical chemical transformations: by dissolving the reaction zone, they facilitate the supply of oxygen to lipids, integrating into lipid complexes, they cause absorption of radiation by them, and reduce the quenching of excited emitter molecules in the medium.

Considering the quantum chemical requirements for radiation production, it can be assumed that inert gases affect the acceleration of VT relaxation and contribute to the effective emptying of lower energy levels (*g*′′ *v*′′), increasing the VT relaxation rate, light inert gases reduce the quenching of emitter molecules on the medium, and, accordingly, the transfer of energy to other molecules, and reduce the radiative decay time of the excited state, contributing to their effective energy loss in the form of radiation, and thus most likely leading to the breakdown of the oxidation chain, creating inactive products. Accordingly, the more severe the hyperlipidemia caused by the disease, the longer the effect of inert gases on the body should be.

Thus, light inert gases, despite their chemical passivity, show significant potential for use in medicine by affecting free radical oxidation.

